# The role of isoforms in the evolution of cryptic coloration in *Peromyscus* mice

**DOI:** 10.1101/041087

**Authors:** Ricardo Mallarino, Tess A. Linden, Catherine R. Linnen, Hopi E. Hoekstra

## Abstract

A central goal of evolutionary biology is to understand the molecular mechanisms underlying phenotypic adaptation. While the contribution of protein-coding and *cis*-regulatory mutations to adaptive traits have been well documented, additional sources of variation—such as the production of alternative RNA transcripts from a single gene, or isoforms—have been understudied. Here, we focus on the pigmentation gene *Agouti*, known to express multiple alternative transcripts, to investigate the role of isoform usage in the evolution of cryptic color phenotypes in deer mice (genus *Peromyscus*). We first characterize the *Agouti* isoforms expressed in the *Peromyscus* skin and find two novel isoforms not previously identified in *Mus*. Next, we show that a locally adapted light-colored population of *P. maniculatus* living on the Nebraska Sand Hills shows an up-regulation of a single *Agouti* isoform, termed 1C, compared to their ancestral dark-colored conspecifics. Using *in vitro* assays, we show that this preference for isoform 1C may be driven by isoform-specific differences in translation. In addition, using an admixed population of wild-caught mice, we find that variation in overall *Agouti* expression maps to a region near exon 1C, which also has patterns of nucleotide variation consistent with strong positive selection. Finally, we show that the independent evolution of cryptic light pigmentation in a different species, *P. polionotus*, has been driven by a preference for the same *Agouti* isoform. Together, these findings present an example of the role of alternative transcript processing in adaptation and demonstrate molecular convergence at the level of isoform regulation.

## Introduction

Understanding the molecular basis of adaptation is one of the principal goals of evolutionary biology. Considerable efforts have been focused on the contributions of protein-coding and *cis*-regulatory mutations, and their relative importance, to phenotypic change (Carroll 2005; Hoekstra & Coyne 2007; Stern & Orgogozo 2008). In contrast, the production of multiple isoforms through the inclusion of different exons in mRNA—known as alternative mRNA processing—is a major source of genetic variation whose role in phenotypic adaptation has been comparatively understudied. A single protein-coding gene can produce multiple isoforms either through alternative splicing, which results in transcripts with different combinations of exons, or through the use of alternative transcription initiation or termination sites, which generate mRNAs that differ at the 5’ or 3’ untranslated regions (UTRs). Alternative processing is ubiquitous throughout eukaryotic evolution but is most prevalent in higher eukaryotes, with mammals having the highest genome-wide rate of alternative splicing events (Barbosa-Morais *et al*. 2012; Merkin *et al*. 2012) and alternative promoter usage (Landry *et al*. 2003; Baek *et al*. 2007; Shabalina *et al*. 2014). In addition, recent studies of mammalian gene expression show that transcripts from the majority of protein-coding genes, especially in primates, undergo alternative processing and generate different isoforms (Blencowe 2006; Kim *et al*. 2008). Thus, by increasing transcriptomic, and hence proteomic diversity, the production of multiple isoforms through alternative processing has been widely regarded as a key mechanism for generating phenotypic diversity and organismic complexity, particularly at large taxonomic scales (Barbosa-Morais *et al*. 2012; Merkin *et al*. 2012).

To investigate the role of alternative processing in adaptation between closely related species, we studied the pigmentation locus *Agouti*, which has been used as a model for isoform régulation in development (Vrieling *et al*. 1994) and has been repeatedly implicated in the evolution of natural variation in pigmentation (e.g., Rieder *et al*. 2001; Schmutz & Berryere 2007; Seo *et al*. 2007; Steiner *et al*. 2007; Linnen *et al*. 2009, 2013). *Agouti* encodes a secreted paracrine factor that induces pigment-producing cells (melanocytes) in hair follicles to switch from the synthesis of black pigment (eumelanin) to yellow pigment (phaeomelanin) during hair growth (Jackson 1994). Experiments performed in laboratory mouse (*Mus musculus*) neonates found that the *Agouti* locus comprises three constitutively transcribed coding exons and four upstream, alternatively transcribed 5’ UTRs, also called non-coding exons (Vrieling *et al*. 1994). Thus, in *M. musculus*, *Agouti* mRNAs are found as four different isoforms, each containing one to two non-coding exons upstream of the coding sequence. The “ventral-specific” non-coding exons 1A and 1A’ are expressed in the ventral mesenchyme during embryonic development, whereas the “hair-cycle specific” non-coding exons 1B and 1C are expressed during hair growth across all regions of the body (Vrieling *et al*. 1994). Thus, alternative promoters of *Agouti* allow for differences in the spatial and temporal deployment of a single coding sequence.

Deer mice (genus *Peromyscus*) populations vary tremendously in both color and pattern. Across populations, there is often a close correspondence between the color of mouse fur and the local soil, suggesting that color-matching is important for survival (Dice 1940; 1941; Haldane 1948). In both Nebraska and Florida, dark-colored mice have colonized extreme light substrate environments that appeared in the last 10,000 years— the Sand Hills in Nebraska (Ahlbrandt & Fryberger 1980; Loope & Swinehart 2000) and the coastal islands in Florida (Campbell 1985; Stapor & Mathews 1991). In both cases, mice have independently evolved significantly lighter coats than mice inhabiting darker surrounding substrates (Fig. 1A and 1B), and there is experimental evidence that visually hunting predators generate strong selection favoring substrate matching (Dice 1941; Vignieri *et al*. 2010; Linnen *et al*. 2013).

**Figure 1.**
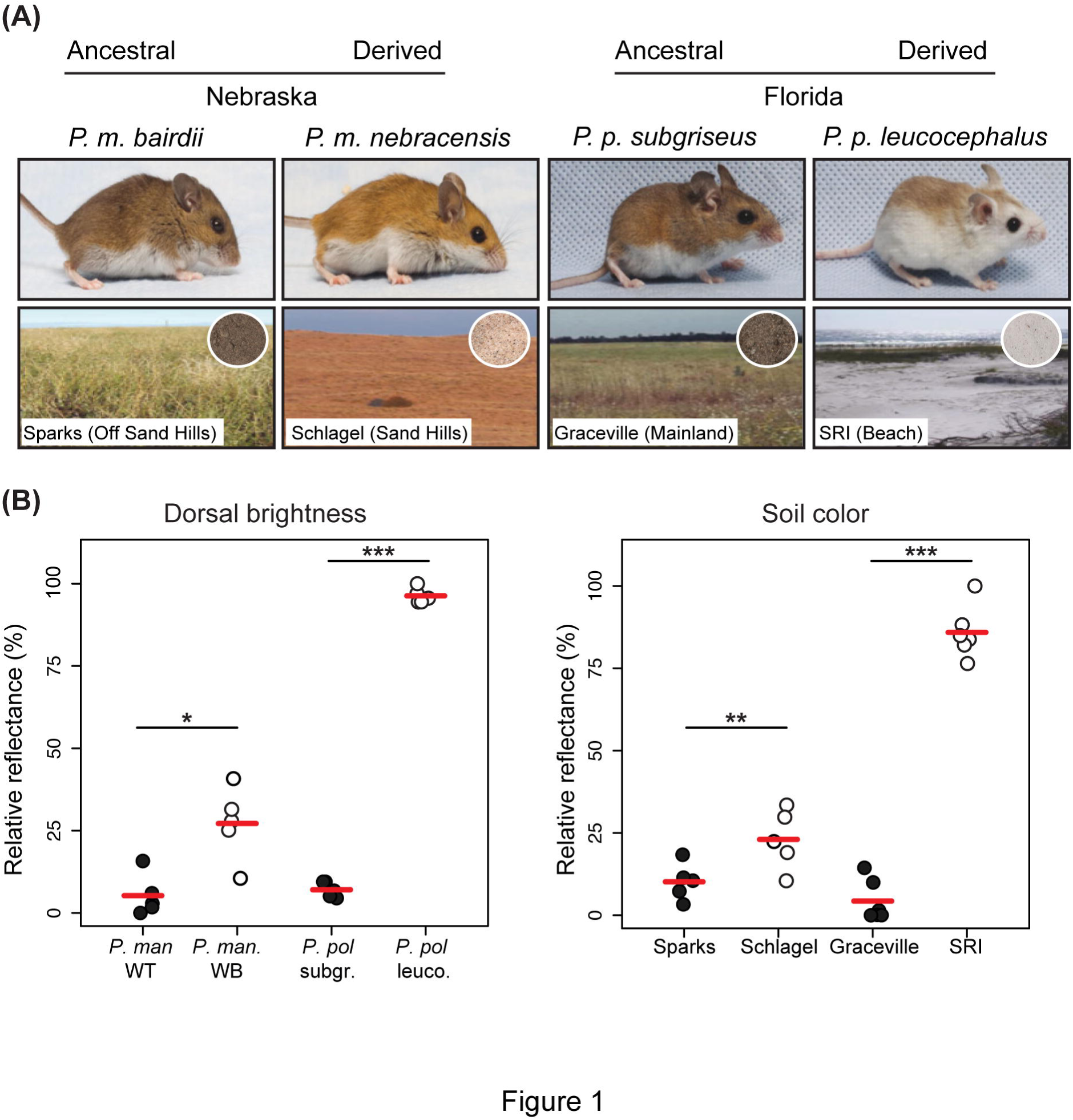
Environmental and phenotypic differences between ancestral and derived habitats and mice in Nebraska and Florida. (**A**) Photographs of wild-type *Peromyscus maniculatus bairdii* (WT), wideband *P*. *m*. *nebrascensis* (WB), *P. polionotus subgriseus* mainland, and *P*.*p. leucocephalus* beach mice (top), typical habitats (bottom), and soil (bottom inset). (**B**) Dorsal brightness (total dorsal reflectance) measured in mice from ancestral (black circles) and derived (white circles) habitats. (**C**) Reflectance of soil samples from ancestral (black circles) and derived (white circles) habitats. Soil samples are from Cherry County, Sparks, Nebraska vs. Cherry County, Schlagel, Nebraska; Jackson County, Graceville, Florida vs. Okaloosa County, Santa Rosa Island (SRI) Florida. * P < 0.05, **P < 0.01, ***P < 0.001, two-tailed *t*-tests; *n* = 5 (mice) and *n* = 5 (soil). Reflectance values were normalized by subtracting the darkest value and dividing by the lightest minus the darkest.

In addition to phenotypic convergence, the same gene, *Agouti*, has been shown to play a significant role in mediating this color adaptation in both Nebraska and Florida populations (Steiner *et al*. 2007; Linnen *et al*. 2009; Manceau *et al*. 2011; Linnen *et al*. 2013). Specifically, in light-colored *P. maniculatus* inhabiting the Nebraska Sand Hills, referred to as “wideband” mice, *Agouti* is expressed at higher levels and for a longer period during hair growth than in their ancestral dark-colored counterparts (“wild-type” mice), which gives rise to a lighter and wider pheomelaninic band on individual hairs and overall lighter appearance (Linnen *et al*. 2009). Furthermore, wideband mice display marked differences in additional pigment traits (e.g., dorso-ventral boundary, ventral color, and tail stripe) that are significantly associated with variants in the *Agouti* locus (Linnen *et al*. 2013). In Florida, changes in *Agouti* and two other pigmentation loci are responsible for producing the lighter pigmentation phenotypes displayed by a derived population of beach mice (*P. polionotus leucocephalus*) inhabiting the coastal sand dunes, relative to their ancestral mainland conspecifics (*P*.*p*. *subgriseus*) (Hoekstra *et al*. 2006; Steiner *et al*. 2007; Mullen & Hoekstra 2008). Like wideband mice from the Nebraska Sand Hills,Florida beach mice express *Agouti* at higher levels and in extended spatial domains compared to mainland mice (Manceau *et al*. 2011).

The implication of *Agouti* in the repeated evolution of cryptic coloration in *Peromyscus*, coupled with its regulatory architecture of alternative 5’ non-coding exons expressed in a region-and temporal-specific pattern, make this an excellent system in which to study the role of isoform regulation in evolution and adaptation. Here, we first characterize the *Agouti* isoforms present in *Peromyscus* skin and examine their patterns of expression during hair growth. We then use *in vitro* experiments to study transcript-specific functional différences. Next, we use a phenotypically variable population in the Sand Hills to map variation in overall *Agouti* mRNA expression to the *Agouti* locus and to test for signatures of selection on the different isoforms. Finally, we study the expression of *Agouti* isoforms in Florida *P. polionotus* to establish whether the changes in isoform regulation seen in Nebraska *P. maniculatus* are shared with its sister species that independently evolved light coloration as an adaptation to a similar light-colored sandy environment.

## Materials and Methods

### Mice

#### Lab mice strains

We originally acquired wild-derived strains from the *Peromyscus* Genetic Stock Center (University of South Carolina) and now maintain them at Harvard University. As representatives of the mice found in Nebraska, we used *P. maniculatus* wild-type (*a*^+^/*a*^+^) and *P. maniculatus* heterozygous for the wideband allele and a non-agouti allele (*a*^*wb*^/*a*^−^). The non-Agouti allele has a 125-kb deletion that removes the entire regulatory region and first coding exon, resulting in a complete loss of expression (Kingsley *et al*. 2009); thus, *P. maniculatus a*^*wb*^/*a*^−^ are effectively hemizygous for the dominant wideband allele. For many of the experiments, however, we also generated wideband mice homozygous for the wideband allele (*a*^*wb*^/*a*^*wb*^). As representatives of mice in Florida, we used *P. polionotus subgriseus* mainland mice and *P*.*p. leucocephalus* Santa Rosa Island beach mice. All experiments described here were evaluated and approved by Harvard University’s IACUC committee and performed following their established guidelines and regulations.

#### Wild-caught mice

Mice used for the association and selection studies were collected as described in Linnen *et al*. (2009) from two Sand Hills sites located <15km apart in Cherry County, Nebraska (site 1: Schlagel Creek Wildlife Management Area, N=62; site 2: Ballard’s Marsh Wildlife Management Area, N=29).

### Quantification of phenotype and soil coloration

We prepared flat skins of mice from the strains mentioned above using standard museum protocols, which were then deposited in the Mammal Department of the Museum of Comparative Zoology at Harvard University. For all the specimens, we quantified dorsal coloration using a USB4000 spectrophotometer and a PX-2 pulsed xenon light source, and recorded readings using the program SpectraSuite (Ocean Optics). Using a reflectance probe with a shield cut at a 45° angle (to minimize diffuse reflection), we took and averaged three measurements of the dorsal skin of each mouse. We trimmed the data to 300-700 nm range, which represents the visible spectrum of most visual predators (Bennet & Lamoreux 2003), and extracted seven color summary statistics using the program CLRvars (Montgomerie R, 2008, CLR, version 1.05; available at http://post.queensu.ca/~mont/color/analyze.html): B2 (mean brightness), B3 (intensity), S3 (chroma), S5c (chroma), S6 (contrast, amplitude), H3 (hue), and H4c (hue). After performing a normal quantile transformation on each variable, we performed a principal components analysis (PCA) on the transformed data in R (R: A Language and Environment for Statistical Computing, R Core Team, 2014, http://www.R-project.org/) using prcomp.

To quantify variation in soil color, we collected soil samples from four different localities representing the natural habitats of the mice used in this study: Schlagel Creek Wildlife Management Area, Cherry County, Nebraska (*P. maniculatus* wideband mice); Sparks, Cherry County, Nebraska (*P. maniculatus* wild-type mice); Santa Rosa Island, Okaloosa County, Florida (*P*.*p. leucocephalus* beach mice); and Graceville, Jackson County, Florida (*P*.*p. subgriseus* mainland mice). We quantified soil reflectance with a spectrophotometer as described above.

### Rapid Amplification of cDNA Ends (RACE)

We sampled dorsal and ventral skin of *P. maniculatus* wideband (*a*^*wb*^/*a*^−^) and wild-type strains and used 5’ rapid amplification of cDNA ends (RACE) to identify the *Agouti* isoforms present in *Peromyscus*. To explore and characterize the full diversity of *Agouti* isoforms, we sampled tissue from pups (postnatal day 4 [P4]), as previously done in *Mus* (Vrieling *et al*. 1994), and from adults (>30 days), since the pelage in adult *Peromyscus* differs from juveniles (Fig. S1, supporting information; Golley *et al*. 1966), and therefore may contain additional transcripts not found during the initial hair cycle.

For each mouse strain, we extracted RNA from skin taken from 1-4 individuals. We dissected skin immediately after sacrifice and stored it in RNAlater (Qiagen) at 4^0^C. We extracted total RNA using the Qiagen RNeasy Fibrous Tissue Kit. To maximize tissue disruption and RNA yield, we performed two 2-min homogenizations at 50Hz using a 5mm steel ball and a Tissuelyser LT (Qiagen). Following extraction, we quantified RNA using a Quant-it RNA kit and a Qubit flourometer (Invitrogen). From total RNA, we then purified mRNA using NucleoTrap mRNA kits (Clontech) and quantified mRNA using fluorescence as described above.

Following mRNA purification, we used the SMARTer RACE cDNA Amplification Kit (Clontech) to prepare 5’-RACE ready cDNA and perform RACE PCRs. To increase the specificity of the 5’ RACE PCRs, we performed a nested PCR. For the first RACE PCR, we used the kit-provided UPM primer in conjunction with a primer located in the exon 4 of *Agouti* (“GSP1” sequence: GTTGAGTACGCGGCAGGAGCAGACG). For the nested PCR, we used the kit-provided NUPA primer in conjunction with a primer located upstream of GSP1 (“NGSP1C”, sequence: TCTTCTTCAGTGCCACAATAGAAACAG). Following the second PCR, we gel-purified any obvious bands using a Qiaquick gel extraction kit (Qiagen). We then ligated the resulting DNA to vectors and transformed competent cells using the pGEMt easy Vector kit (Promega).

Following transformation, we performed colony PCRs using Qiagen TAQ and M13F and M13R primers. PCR products were purified enzymatically using Exonuclease I and Shrimp Alkaline Phosphatase (USB). Purified PCR products were sequenced on an ABI 3730xl Genetic Analyzer at Harvard University’s Genomics Core facility. In total, we obtained sequences from 24-89 *Agouti* clones from each of eight distinct genotype/stage/tissue combinations. Then, all *Agouti* clones were sequenced using primers placed at the 5’ end and with *Agouti*-Exon3-R (see Supplementary Table 1) to confirm the presence of the *Agouti* coding sequence.

### Quantitative PCR (qPCR)

#### Lab strains

We performed gene expression analyses in P. maniculatus wideband (*a*^*wb*^/*a*^*wb*^), *P. maniculatus* wild-type, *P*.*p. subgriseus*, and *P*.*p. leucocephalus mice*. In all cases, we used age-matched mice to minimize the variability in *Agouti* expression introduced when hair follicles are at different stages of the hair cycle. We extracted RNA from dorsal and ventral tissue as indicated above, isolated mRNA from total RNA using the NucleoTrap mRNA Kit (Clontech), and directly synthesized cDNA using qScript cDNA SuperMix (Quanta BioSciences). We performed all reactions in triplicate using PerfeCta SYBR Green FastMix (Quanta BioSciences) and calculated relative expression between samples using the 2^—ΔΔC^_T_ method (Livak & Schmittgen 2001), using *β*-actin as a reference gene.

To measure levels of total *Agouti*, we used the primers *Agouti*-Exon2-F and *Agouti*-Exon3-R to amplify all transcripts containing the *Agouti* coding sequence. To measure levels of individual isoforms, we paired a forward primer within the noncoding exon (1C-F, 1D-F, or 1E-F, respectively) with a reverse primer in the coding region (*Agouti*-Exon2-R). We amplified *β*-actin using the primers *β*-actin-Pero-F and *β*-actin-Pero-R. All primer sequences were designed from the *Peromyscus maniculatus* genomic reference (Pman_1.G, GenBank accession: GCA_000500345.1) and can be found in Supplementary Table 1.

#### Wild-caught mice

We removed a 5mm skin biopsy from the dorsum of recently sacrificed mice, preserved tissue in RNAlater, and extracted RNA as indicated above. We synthesized cDNA and performed qPCR reactions to measure total *Agouti* and *β*-actin as described for lab strains.

### Dorsal depilation

It is well established that *Agouti* expression is tightly linked to hair growth (Vrieling *et al*. 1994). Contrary to newborn pups, in which hair emerges simultaneously across the body, hair follicles of adult mice are desynchronized with respect to the hair cycle (Muller-Rover & Paus 2001). Thus, the quantification of *Agouti* expression in adult dorsal skin represents an average expression level across many follicles that are each at different stages in the cycle. To understand the detailed dynamics of *Agouti* isoform expression across the entire hair cycle in adults, we depilated the backs of adult wild-type and wideband mice, a procedure that resets and synchronizes the hair follicle program (Paus & Cotsarelis 1999), and compared isoform expression at different time points. We anesthetized adult *P. maniculatus* wideband and wild-type mice by performing intraperitoneal injections with a cocktail of Ketamin/Xylazine (0.1mL/20g mouse wt) and depilated a small patch (~1cm^2^) in the dorsum using a melted beeswax/resin mixture. After the procedure, we applied topical Lidocaine for pain relief every hour for the next eight hours.

### mRNA stability assays

Our RACE experiment revealed that the dorsal skin in *Peromyscus* expresses three *Agouti* isoforms (1C, 1D, and 1E) simultaneously, differing only in the first, non-coding exon. To determine whether there were differences in mRNA half-life, we cloned each of the three isoforms from *P. maniculatus* wild-type into a pHAGE-CMV-eGFP-W expression vector such that the noncoding exon and *Agouti* coding sequence replaced the eGFP coding sequence. We used Lipofectamine (Life Technologies) to transfect plasmids into human embryonic kidney (HEK293) cells and 40 h after transfection halted transcription by treating cells with 10 μg/ml actinomycin D (Sigma) in DMSO. HEK cells were used in these stability assays because they provide a suitable cellular environment for transcription of mRNA encoded by expression vectors; in addition, they do not express *Agouti* endogenously, so measurements performed (see below) accurately reflect the amount of transcript produced from the transfected expression vector alone. As a control, we treated cells with an equal volume of 100% DMSO. At Gh, 2h, or 4h after actinomycin treatment, we collected cells in TRIzol reagent (Invitrogen) and extracted RNA using the Direct-zol RNA Mini Kit (Zymo Research). We then performed qPCR as described above, using primers common to all constructs (i.e., located in exons 2 and 3): Agouti-Exon2-F and Agouti-Exon3-R to amplify *Agouti* and *β*-actin-Human-F and ‐R to amplify endogenous *β*-actin. Primer sequences can be found in Supplementary Table 1. We used three replicates for each condition and time point. *Agouti* expression in actinomycin-treated cells at each time point was calculated as a percentage of *Agouti* expression in DMSO-treated cells at the same time point using the 2^—ΔΔC^_T_ method. We then tested for significant differences between the half-lives of the isoforms by comparing the slopes of the respective trendlines, using one-way analysis of covariance (ANCOVA).

### Luciferase assays

To establish whether the non-coding exons of the isoforms expressed in the dorsal skin of *Peromyscus* showed differences in translation, we generated luciferase reporter plasmids by cloning each of the three noncoding exons (1C, 1D, or 1E) from both *P. maniculatus* wideband and wild-type strains into a 5’ UTR reporter vector (pLightSwitch_5UTR; Switchgear Genomics) upstream of the luciferase coding sequence and downstream of the ACTB promoter. We generated mutated 1D reporter plasmids from the wild-type 1D plasmid using the Q5 Site-Directed Mutagenesis Kit (New England BioLabs). We transfected plasmids into HEK293 cells using Lipofectamine (Invitrogen), with six replícate transfections per construct. Forty-eight hours after transfection, we measured luminescence as a readout of protein production using a microplate reader (SpectraMax L). We quantified transcription from each plasmid as follows: we collected six replicates of transfected cells in TRIzol reagent (Invitrogen) and extracted RNA using the Direct-zol RNA Mini Kit (Zymo Research). We then carried out qPCR as described above using the primer pairs Luciferase-F and ‐R, and *β*-actin-Human-F and ‐R. Primer sequences can be found in Supplementary Table 1.

### Association tests

Using association mapping in a variable, natural population, previous work has shown that multiple mutations in *Agouti* are statistically associated with different aspects of coat color (Linnen *et al*. 2013). These mutations may affect phenotype via measurable changes to *Agouti* expression level. To determine whether color phenotypes and *Agouti* expression co-localize to the same *Agouti* region(s), we measured *Agouti* expression (overall *Agouti* and isoform 1C) as described above in 88 mice that had been genotyped and scored for color traits as described in Linnen *et al*. (2013). To test for an association between *Agouti* genotype and expression, we performed single-SNP linear regressions in PLINK v1.07 (Purcell *et al*. 2007). To control for population structure, we genotyped individuals at 2,077 unlinked SNPs as described in Linnen *et al*. (2013), and used SMARTPCA to conduct a PCA, followed by TWSTATS to evaluate the statistical significance of each principle component (Patterson *et al*. 2006). We detected four significant genetic principal components and included these as covariates in the association analysis. Prior to analysis, we performed a normal-quantile transformation on the phenotype and expression data in R to ensure that our phenotypic data were normally distributed (Guan & Stephens 2011). To correct for multiple testing, we used the step-up method for controlling the False Discovery Rate, which we set to a threshold of 10% (Benjamini & Hochberg 1995).

## Results

### *Peromyscus* **expresses two** *Agouti* **isoforms not previously identified in** *Mus*

RACE from *P. maniculatus* ventral samples revealed the presence of the ventral specific isoforms 1A, 1A’, and 1A1A’ (an isoform containing both the 1A and 1A’ exons), and the hair-cycle specific 1C, similar to what is seen in *M. musculus* (Vrieling *et al*. 1994). However, none of our clones contained sequences corresponding to exon 1B found in *M. musculus* (Vrieling *et al*. 1994) (Fig. 2A and Fig. S2, supporting information). Our RACE experiment in dorsal skin revealed that, in addition to hair-cycle specific isoform 1C, *P. maniculatus* expressed two additional isoforms of *Agouti*, hereafter referred to as 1D and 1E, which had not been previously reported in M. musculus (Fig. 2A,B and Fig. S2, supporting information). Like the previously described *Agouti* isoforms, 1D and 1E each consist of an alternate non-coding exon (exon 1D or 1E), followed by three protein-coding exons (exons 2, 3, and 4), which are shared by all *Agouti* isoforms. Both exons 1D and 1E are located downstream of exons 1A and 1C in the *Agouti* locus (Fig. 2B) and show polymorphisms between wideband and wild-type *P. maniculatus* (Fig. S2, supporting information). To confirm the absence of isoform 1B in ventral/dorsal tissue and of isoforms 1D and 1E in ventral tissue, as indicated by our RACE experiments, we carried out qPCR using isoform-specific primers in the respective tissues and did not detect any expression of these transcripts. Together, our results indicate that *P. maniculatus* expresses some, but not all, of the *Agouti* isoforms previously described in *M. musculus* (the ventral-specific 1A, 1A’, 1A1A’ and the hair cycle specific 1C), and also contains two novel dorsal-specific isoforms that had not been described previously (1D and 1E).

**Figure 2.**
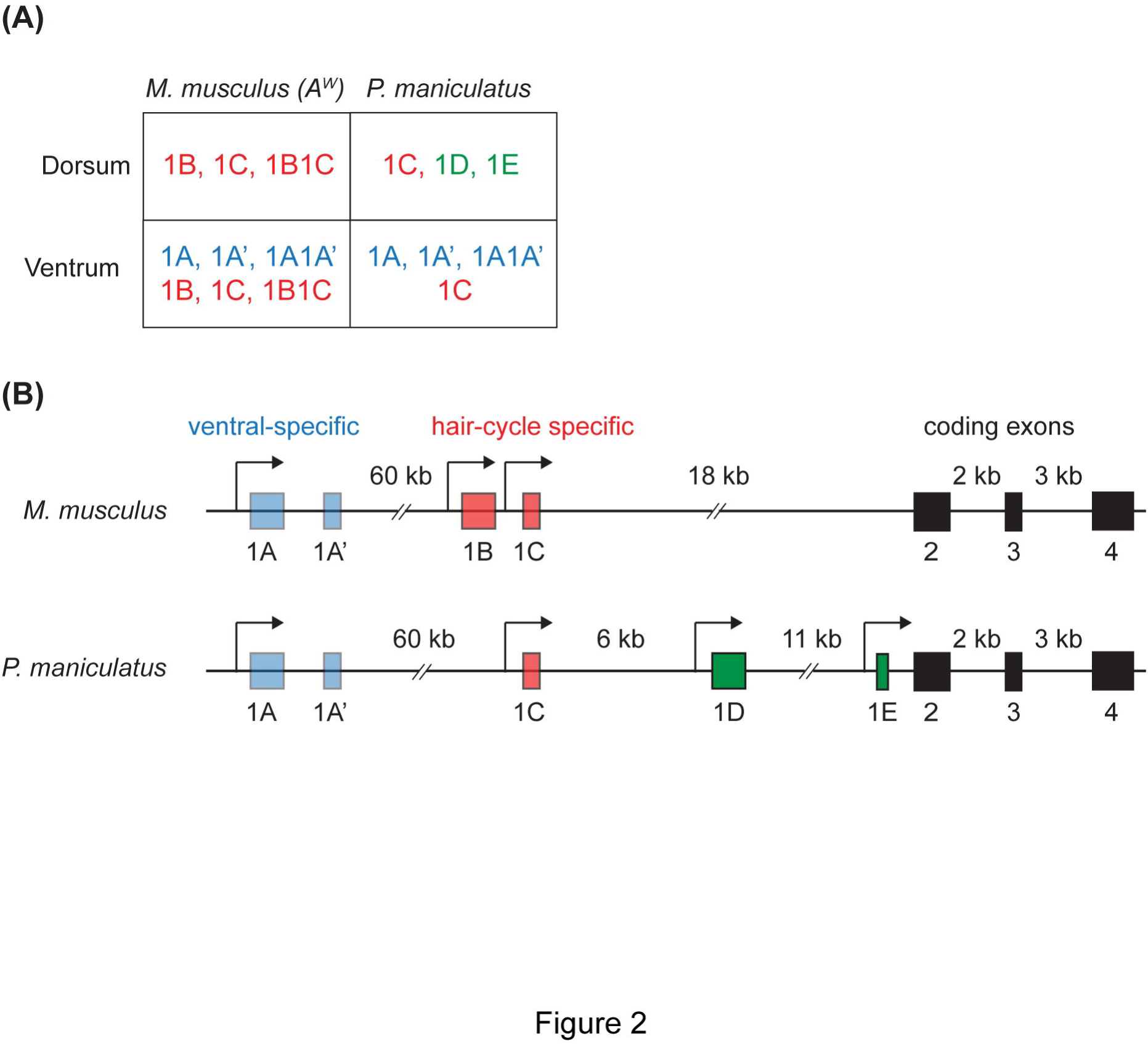
Characterization of *Agouti* isoform expression in *P. maniculatus* using 5’ RACE. (**A**) *Agouti* isoforms expressed in the ventrum and dorsum of *M. musculus* pups (strain A^W^) and wild-type *P. maniculatus*. For *P. maniculatus*, all isoforms were found in both adults and pups by RACE and/or qPCR. (**B**) Map of the *Agouti* locus in *M. musculus* (above) and *P. maniculatus* (below). Colors in (**A**) and (**B**) represent ventral specific non-coding exons (blue), hair cycle-specific non-coding exons (red), and the novel non-coding exons (green) reported here. Coding exons (black) are common to all isoforms.

### **Differences in** *Agouti* **mRNA levels between wideband and wild-type mice are driven by upregulation of isoform 1C**

Wideband and wild-type mice differ markedly in their dorsal coloration; therefore, we focused here on analyzing the isoforms present in the dorsum. Our RACE experiments demonstrated that the dorsal skin of *P. maniculatus* expresses at least three different *Agouti* transcripts (1C, 1D, and 1E) simultaneously, but it remained unknown whether all three contribute to the increase in *Agouti* mRNA levels seen in wideband mice relative to wild-type ones or whether this difference is driven only by a subset. To answer this question, we used qPCR to measure the relative expression of total *Agouti* mRNA and each of its isoforms in dorsal skin of wideband and wild-type *P. maniculatus*. The expression of overall *Agouti* was approximately 22-fold higher in wideband mice than in wild-type mice (*P* = 0.0011, two-tailed t-test) (Fig. 3A). We then measured isoform-specific expression and found that isoform 1C was significantly higher in wideband than in wild-type mice (*P* = 0.0016, two-tailed *t*-test) (Fig. 3B). In contrast, we did not observe significant differences in mRNA levels between the two strains in expression of isoforms 1D or 1E (*P* = 0.3080 and *P* = 0.6286, respectively; two-tailed *t*-tests) (Fig. 3B).

**Figure 3.**
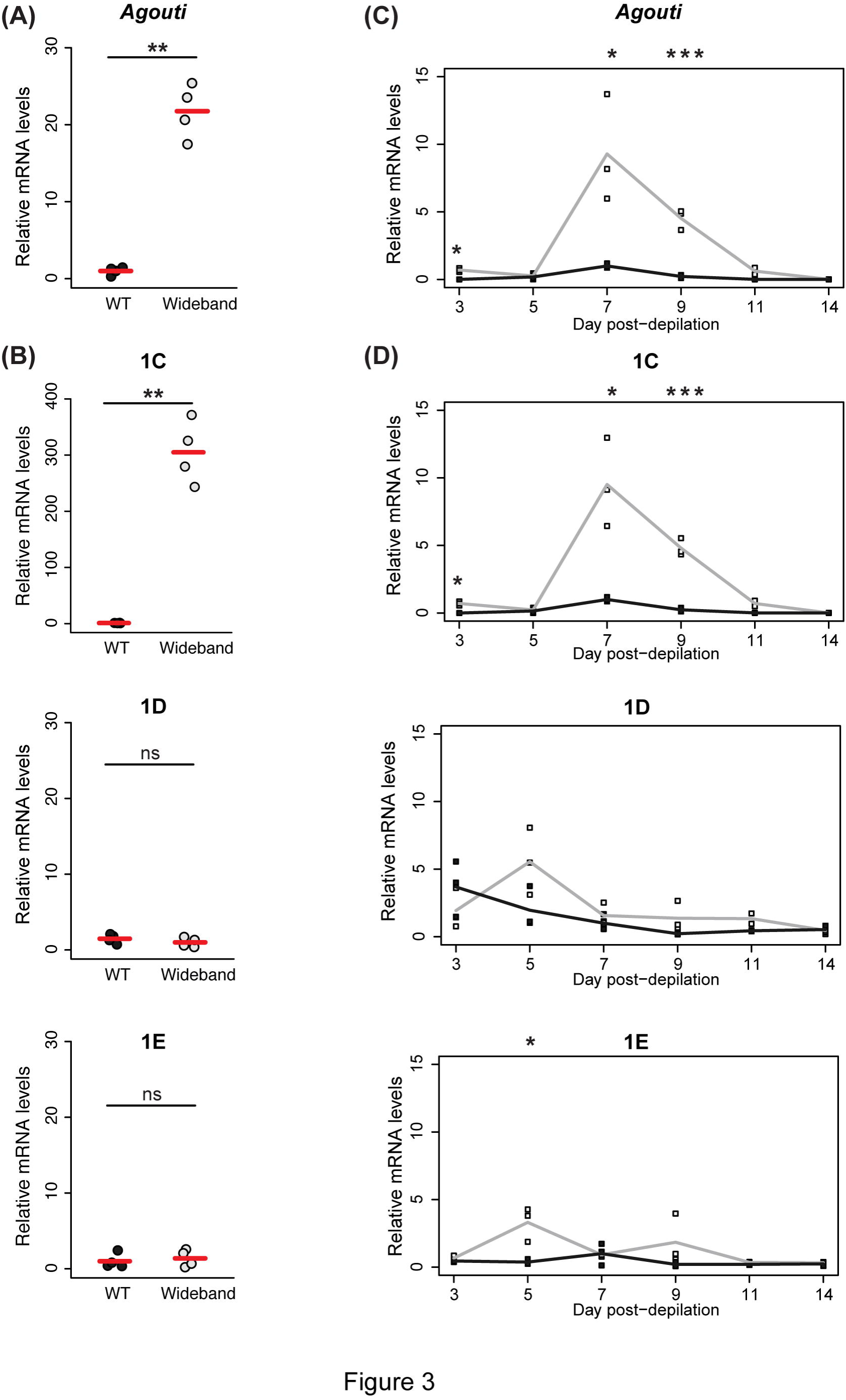
Differential expression of *Agouti* isoforms between wideband and wild-type mice. (**A**) qPCR in adults shows that the overall expression of *Agouti* mRNA in wideband mice (light circles) is higher than wild-type mice (dark circles). (**B**) Isoform-specific mRNA levels of isoform 1C are higher in wideband than wild-type mice, but no significant differences in 1D or 1E expression. (**C**) qPCR measurements of overall *Agouti* mRNA levels at different time points after hair removal show that wideband and wild-type mice differ in the expression of total *Agouti* at days 3, 7, and 9 after depilation. (**D**) Isoform specific measurements show that isoform 1C differs between the two strains at days 3, 7, and 9, whereas isoforms 1D and 1E do not differ between the two strains, with the exception of 1E, which differs at day 5. * P < 0.05, **P < 0.01, ***P < 0.001, two-tailed *t*-tests; *n* = 4 (for each strain in (**A**) and (**B**)) and *n* = 3 (for each strain and time point in (**C**) and (**D**)); red bars indicate the mean.

We next quantified total *Agouti* and isoform-specific expression at different time points following dorsal depilation in order to examine the dynamics of *Agouti* isoform expression during hair growth. In both strains, total *Agouti* is expressed initially at low levels, peaks at day 7 after depilation, and then decreases (Fig. 3C), a pattern that mirrors the expression of *Agouti* during the first hair cycle in pups (Linnen *et al*. 2009). Total *Agouti* levels in wideband mice were higher than in wild-type mice at days 3, 7, and 9 after depilation (*P* = 0.0011, *P* = 0.0228, and *P* = 0.0006, respectively, two-tailed *t*-tests) (Fig. 3C). When we measured transcript-specific mRNA levels in wideband and wild-type mice, we found that the expression of isoform 1C closely matched the expression of overall *Agouti*, peaking at day 7, and differing from wild-type mice also at days 3, 7, and 9 (*P* = 0.0012, *P* = 0.0109, and *P* = 0.0003, respectively, two-tailed *t*-tests) (Fig. 3D). This marked similarity suggests that isoform 1C levels are primarily responsible for explaining overall *Agouti* levels. In contrast, isoform 1D and 1E did not differ between wideband and wild-type mice at any time points, with one exception: expression of 1E was higher in wideband than wild-type at day 5 (*P* = 0.0166, two-tailed *t*-test) (Fig. 3D).

To determine whether the same patterns occur in neonates, we measured isoform-specific mRNA levels at postnatal day 4, which corresponds to the stage in the first hair cycle when *Agouti* expression is highest (Linnen *et al*. 2009). Quantitative PCR confirmed that wideband mice express higher levels of total *Agouti* mRNA relative to wild-type mice (*P* = 0.0010, two-tailed *t*-test) (Fig. S3). In addition, like in adults, isoform 1C expression was significantly higher in wideband mice than in wild-type (*P* = 0.0003, two-tailed *t*-test), whereas there were no significant differences between the two strains in the expression of isoforms 1D or 1E (*P*= 0.1575 and *P* = 0.3231, respectively, two-tailed *t*-tests) (Fig. S3). Together, the results of our measurements of isoform-specific *Agouti* mRNA levels in adults and pups indicate that the marked increase in *Agouti* expression seen in wideband mice, relative to wild-type mice, is primarily driven by upregulation of isoform 1C.

### *Agouti* **isoforms differ in luciferase production**

To investigate functional variation associated with different *Agouti* isoforms, we measured their half-lives *in vitro*. After halting transcription from a vector expressing each isoform, we used qPCR to measure mRNA levels at different time points during the transcript decay that followed and did not detect any statistically significant differences between the half-lives of the three isoforms (*P* = 0.1230, one-way ANCOVA) (Fig. S4). Thus, our experiment indicates that the three alternative exons simultaneously expressed in *Peromyscus* dorsal skin do not have measurable effects on the stability of the *Agouti* transcripts.

To determine if the isoforms differed in their regulation of translation, we next examined whether each of the alternative exons affects the amount of protein produced from a luciferase transcript. Relative to a control vector expressing a luciferase coding sequence without a 5’UTR, a vector carrying exon 1C showed a marked increase in luciferase activity (*P* = 0.0390 [wideband sequence] and *P* = 0.0001 [wild-type sequence], two-tailed *t*-tests), whereas 1D showed a marked decrease (*P* = 4.4 x 10^−5^ [wideband sequence] and *P* = 0.0002 [wild-type sequence], two-tailed t-tests). Exon 1E from wideband mice was not significantly different from the control (*P* = 0.2577 [wideband sequence], two-tailed *t*-test), whereas exon 1E from wild-type mice showed increased luciferase activity compared to the control (*P* = 0.0082, two-tailed *t*-test) (Fig. 4A). Importantly, we found that mRNA levels did not differ between any of the vectors (*P* = 0.3790, ANOVA), demonstrating that differences in translation, not transcription, are exclusively responsible for the differences in luminescence (Fig. 4B).

**Figure 4.**
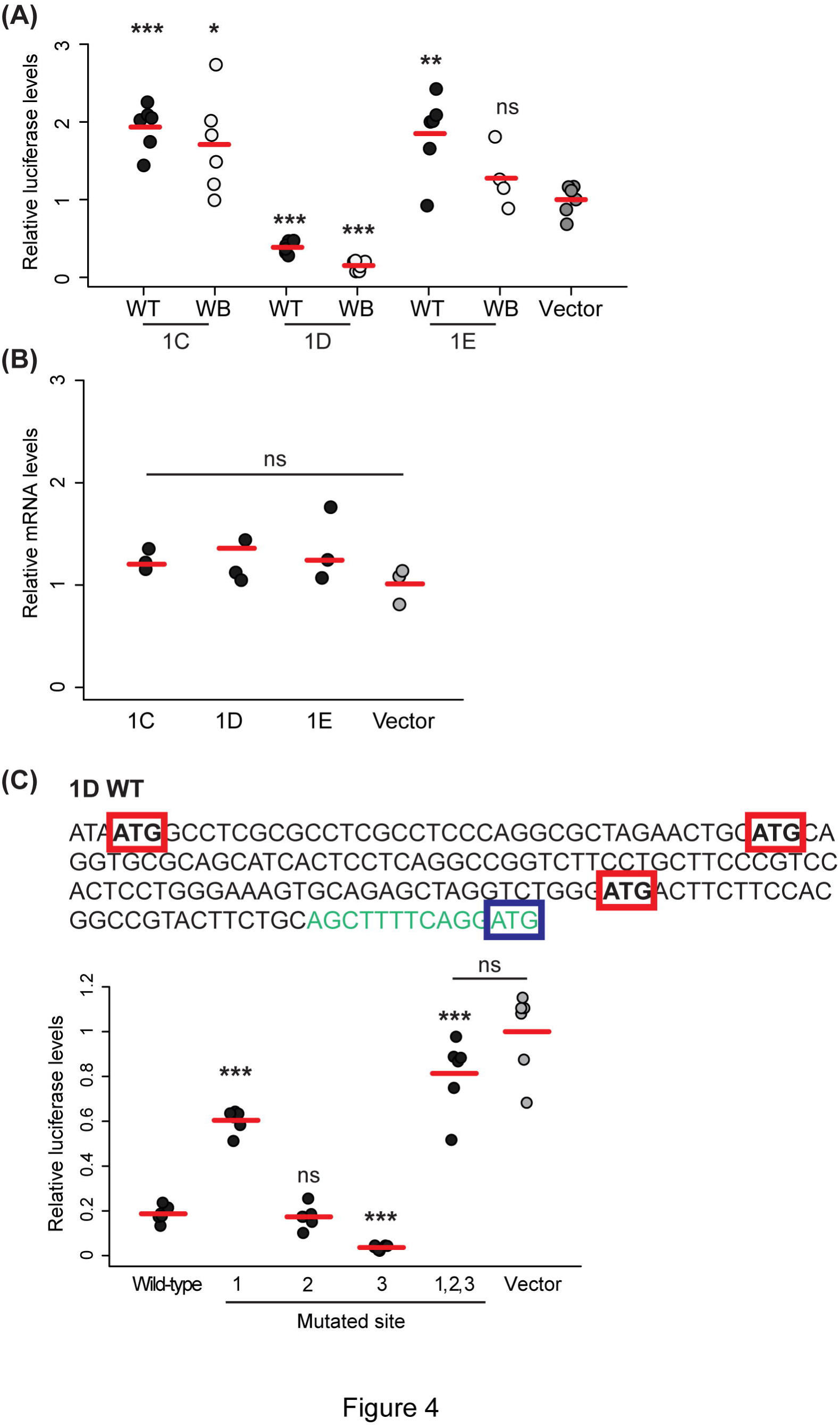
*Agouti* isoforms differ in luciferase production. (**A**) Relative luciferase levels produced by transcripts carrying the three different 5’ UTRs expressed in *P. maniculatus*. * P < 0.05, **P < 0.01, ***P < 0.001, two tailed *t*-tests; n = 6 per construct. (**B**) mRNA levels of luciferase did not differ between the constructs demonstrating that differences seen in (**A**) occur at the posttranscriptional level; ANOVA, *n* = 3. Only results from wild-type sequences are shown. (**C**) (Above) The sequence of exon 1D (black font) in wild-type *P. maniculatus*.Upstream start codons are boxed in red, the beginning of exon 2 is in green font, and the functional ATG is boxed in blue. (Below) Quantification of relative luciferase levels from transcripts carrying the *P. maniculatus* wild-type exon 1D and those carrying a version of exon 1D where each upstream ATG site has been mutated to ACG, relative to a control lacking a 5’ UTR. * P < 0.05, **P < 0.01, ***P < 0.001, two tailed *t*-tests; *n* = 6 per construct; red bars indicate the mean. In all cases, luciferase levels are normalized relative to background levels.

To further dissect the mechanisms underlying the differences in protein translation observed between the isoforms, we examined their sequences in more detail and found that exon 1D contains start codons (ATGs) upstream of the *Agouti* start codon (three in the wild-type 1D sequence and two in the wideband sequence) (Fig. 4C and Fig. S2). Upstream start codons have been shown to decrease translation efficiency by recruiting ribosomes away from the start codon (Kozak 2002; Rosenstiel *et al*. 2007; Song *et al*. 2007; Medenbach *et al*. 2011). To determine whether this can explain the reduced translation of isoform 1D, we mutated the ATG sites to ACG in the *P. maniculatus* wild-type sequence and quantified the relative amount of luciferase produced (Fig. 4C). When we mutated and tested each ATG site individually, we found that mutating site 1 led to a significant increase in luciferase production relative to the wild-type 1D sequence (*P*=1.268 x 10^−8^, two-tailed *t*-test), mutating site 2 resulted in no significant difference(*P* = 0.5966, two-tailed *t*-test), and mutating site 3 led to a significant reduction (*P*=1.6739 x 10^−6^, two-tailed *t*-test) (Fig. 4C). Thus, the different ATG sites in exon 1D differ in their ability to modulate luciferase production, possibly due to differences in their surrounding sequences (Kozak 1999; 2002). When we mutated the three ATG sites simultaneously, luciferase production from the mutant transcript was significantly higher than that from the wild-type 1D transcript (*P* = 0.000015, two-tailed *t*-test) and did not differ from the control (*P* = 0.0920, two-tailed *t*-test), indicating that the upstream start codons contained within exon 1D are responsible for the marked decrease in translation from this transcript. Together, these experiments demonstrate that the non-coding exons of the *Agouti* isoforms expressed in the dorsal skin of *P. maniculatus* differ in their regulation of protein translation, with non-coding exon 1C generating the highest level of luciferase, relative to the control, and non-coding exon 1D with its multiple ATG sites generating the lowest.

### **Genetic variation near exon 1C is associated with dorsal color and** *Agouti* **expression in wild-caught mice**

To evaluate evidence for exon 1C’s contribution to adaptive coat color variation in natural populations, we examined patterns of genotype-color associations (from Linnen *et al*. 2013) and genotype-expression associations (this study) across the 180-kb *Agouti* locus in a phenotypically variable population of *P. maniculatus*. Here, we focus on dorsal brightness because we expect that this trait (1) has a large impact on substrate matching (hence, fitness), and (2) has the potential to be strongly influenced by the hair-cycle isoform 1C due to *Agouti*‐s direct impact on pigment deposition in hairs. For dorsal brightness, we previously identified a peak in association centered directly on exon 1C (Fig. 5A). Intriguingly, we also found a peak in association with expression at the same location (Fig. 5B). We note, however, that these expression data were generated using an assay that detects all *Agouti* isoforms. Nevertheless, when we evaluated the correlation between genotype and isoform 1C expression using 1C-specific probes, the exon 1C SNPs remained significantly associated (SNP 109,902, *P* = 0.0026; SNP 109,882, *P* = 0.0110). Although the lack of polymorphic positions in exon 1C that are derived in wideband mice, relative to the *P. m. rufinus* outgroup (Linnen *et al*. 2009), suggests that the causal mutation is not in exon 1C itself, these association mapping data strongly suggest that there is a causal mutation somewhere in its immediate vicinity that simultaneously increases both expression of isoform 1C and, as a consequence, dorsal brightness.

**Figure 5.**
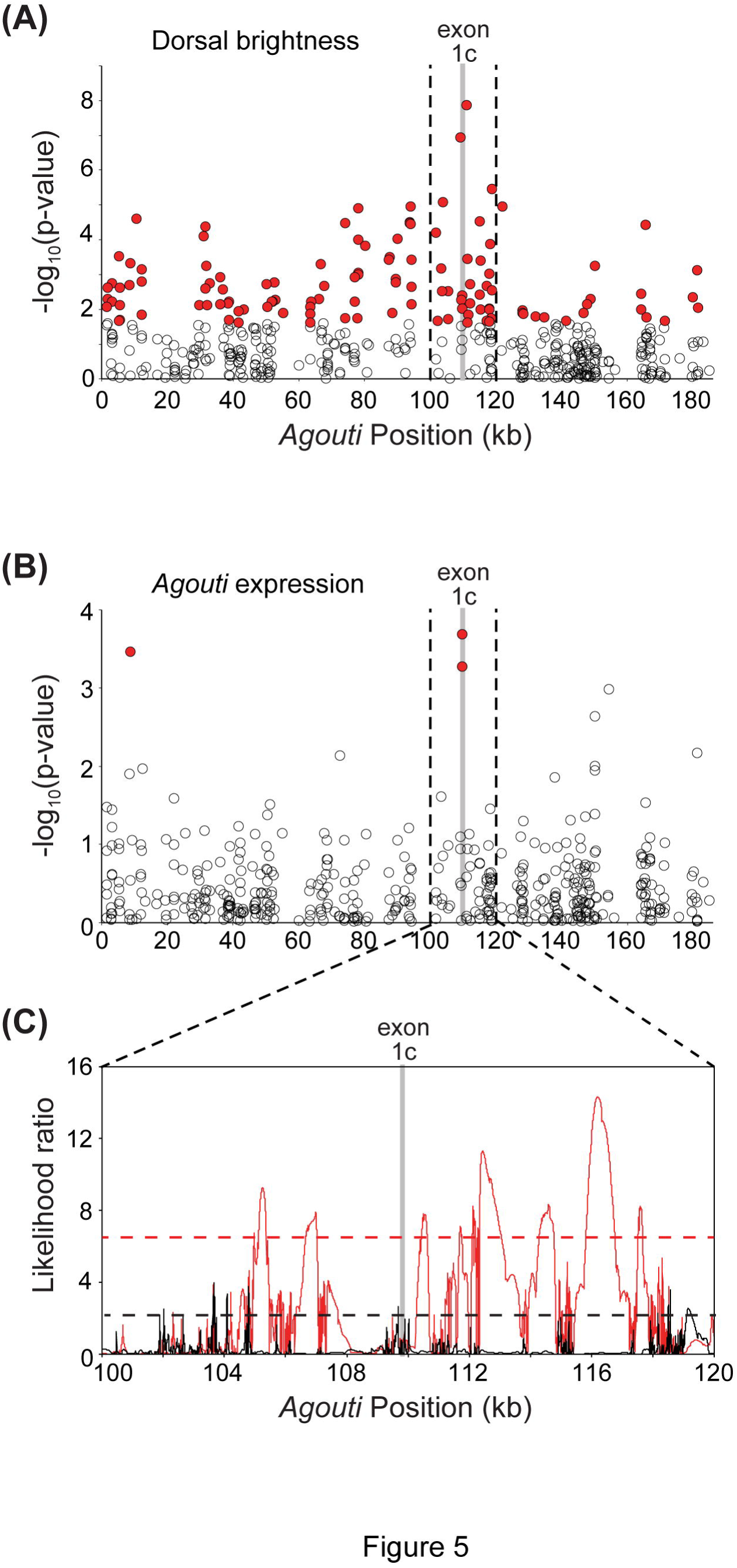
Genetic variation near exon 1C is associated with dorsal color, *Agouti* expression, and signatures of selection in a natural population of *P. maniculatus*. (**A, B**) Strength of statistical association (-log p-value) between dorsal brightness (**A**) and *Agouti* expression (**B**) for 466 SNPs (circles) tested in *n* = 91 (**A**) or 88 (**B**) mice. SNPs significant after a 10% FDR correction are indicated in red. The gray bar highlights the location of exon 1C, and the dashed lines indicate a 20-kb region centered on this exon. Color (**A**) and expression (**B**) both have multi-SNP peaks of association centered on exon 1C. (**C**) Strength of evidence favoring a selection model over a neutral model (likelihood ratios) as a function of location in *Agouti*. Likelihood surfaces are shown for light (red) and dark (black) haplotypes, as determined by the strongest associated SNP in (**A**). Dashed lines indicate significance thresholds, determined via neutral simulations, for light (red) and dark (black) haplotypes. Data for (**A**) and (**C**) are from Linnen *et al*. (2013). *Agouti* positions are defined relative to the *P. maniculatus* BAC clone reported in Kingsley *et al*. (2009).

### Exon 1C has undergone strong positive selection

Although we have not yet identified the causal mutation, the results of our association mapping indicate that it should be in strong linkage disequilibrium with variants located near exon 1C (Fig. 5A, B). In this way, we can use the association mapping results to define light and dark haplotypes (i.e., those that contain the causal mutation and those that do not). If the light mutation has undergone positive selection in the light Sand Hills habitat, we expect to see signatures of selection on the light, but not dark, haplotypes. To evaluate evidence of selection on light and dark haplotypes, we previously (Linnen *et al*. 2013) used the composite-likelihood method implemented in Sweepfinder (Nielsen *et al*. 2005), a method that compares, for each location, the likelihood of the data under a selective sweep to the likelihood under no sweep. The significance of the CLR test statistic is then determined via neutral simulations. In our case, neutral simulations were conducted under a demographic model estimated from genome-wide SNPs (as described in Linnen *et al*. [2013]). Figure 5C depicts the resulting likelihood surfaces and significance thresholds for light and dark haplotypes across a ~20-kb window centered on exon 1C (Fig. 5A, 5B). This analysis indicates strong evidence of selection on the light, but not dark, haplotypes in this region of the *Agouti* locus. Specifically, within this window, the light haplotypes have a 3.6-fold increase in the number of sites rejecting neutrality and a 5.0-fold increase in the average selection coefficient. Together with the results of our association mapping, these analyses indicate that a mutation(s) in the immediate vicinity of exon 1C contributes to dorsal color and is currently undergoing strong positive selection (*s* = 0.14; Linnen *et al*. [2013]).

### **Evolutionary convergence of isoform regulation in** *Peromyscus*

Our results show that *Agouti* isoform 1C is specifically upregulated in the light-colored *P. maniculatus* wideband mice from the Nebraska Sand Hills relative to the dark-colored ancestral population. We next investigated whether similar regulatory mechanisms are observed in another population of *Peromyscus* that independently underwent selection for light pigmentation. We examined patterns of isoform expression in light-colored Santa Rosa Island beach mice (P._*p·*_ leucocephalus) and compared them to dark mainland *P*.^*p·*^ *subgriseus* (Fig. 1). Quantitative PCR revealed that expression of *Agouti* was approximately three-fold higher in beach mice compared to mainland mice (*P* = 0.0142, two-tailed *t*-test; Fig. 6A). Measurements of *Agouti* isoform-specific mRNA levels revealed that there were no significant differences in the expression of isoform 1D or 1E between beach mice and mainland mice (*P* = 0.5352 and *P* = 0.3826, respectively, two-tailed *t*-test; Fig. 6B). In contrast, we found that isoform 1C was significantly upregulated in beach mice compared to mainland mice (*P* = 0.0011, two-tailed *t*-test; Fig. 6B). Together, our measurements of mRNA levels demonstrate that the increase in *Agouti* expression seen in beach mice, relative to mainland mice, is produced primarily by a specific upregulation of isoform 1C, a pattern that matches what we observed in *P. maniculatus* (Fig. 3A).

**Figure 6.**
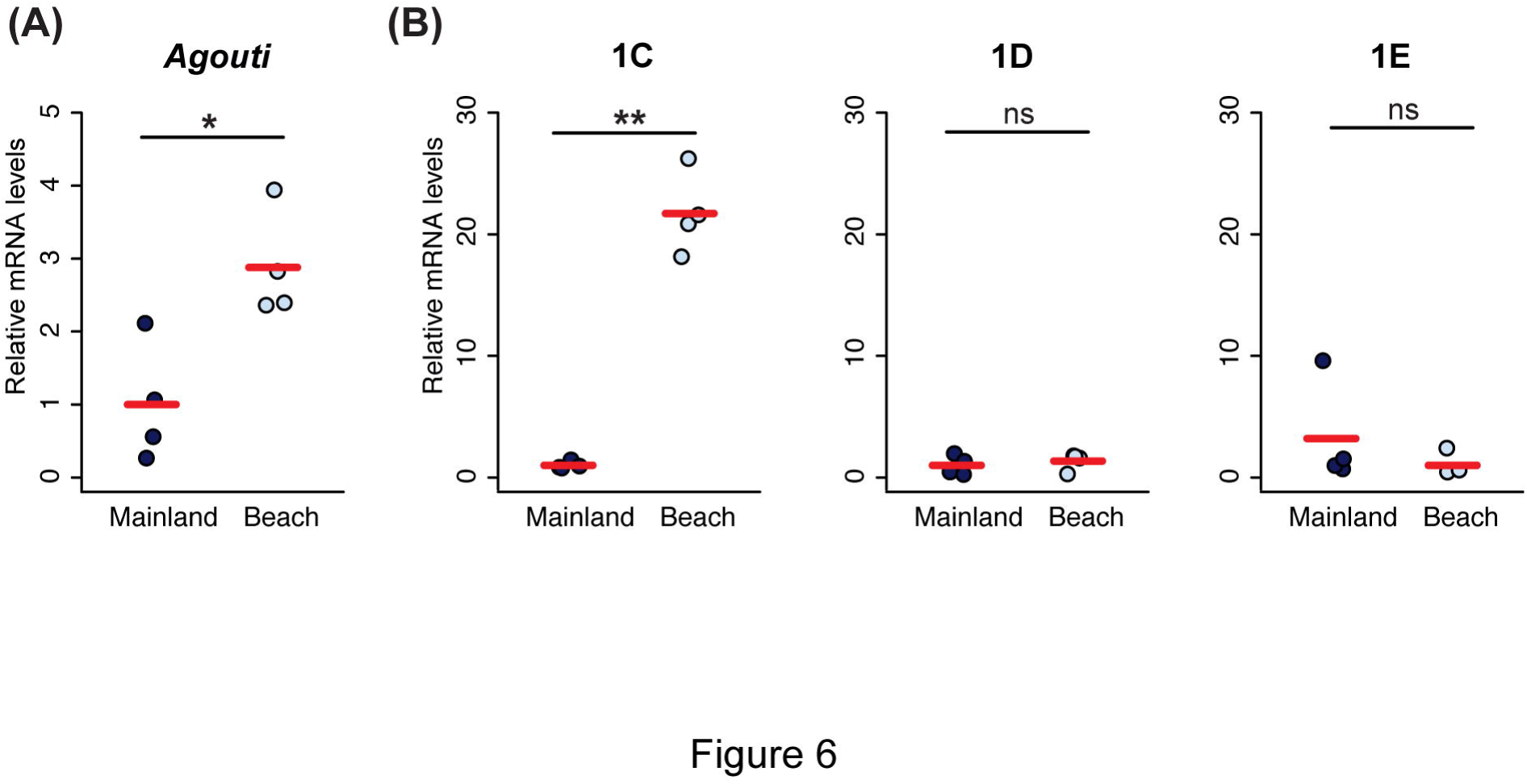
Differential expression of *Agouti* isoforms in *P. polionotus* mainland and beach mice. (**A**) Beach mice (light circles) have higher expression of total *Agouti* than mainland mice (dark circles), as determined by qPCR. (**B**) Expression of isoform 1C is higher in beach mice (light circles) compared to mainland mice (dark circles), whereas there were no significant differences in 1D or 1E expression. * P < 0.05, **P < 0.01, ***P < 0.001, two-tailed *t*-tests; *n* = 4; red bars indicate the mean.

## Discussion

The *Agouti* locus, which contains multiple independently regulated transcription start sites and has been linked to pigment variation in *Peromycus* (Steiner *et al*. 2007; Mullen & Hoekstra 2008; Linnen *et al*. 2009; 2013) and other mammals (Rieder *et al*. 2001; Schmutz & Berryere 2007; Seo *et al*. 2007), represents an ideal study system to understand the importance of alternative transcript processing in adaptation to new environments. While different isoforms have been well studied in *Mus musculus*, in *Peromyscus* we both identify new dorsally expressed isoforms (1D and 1E) as well as the lack of expression of 1B across the body. These differences are consistent with genome-wide surveys of isoform variation, which find rapid evolution of isoform usage between species (Barbosa-Morais *et al*. 2012; Merkin *et al*. 2012).

In this study, we also find that although the dorsal skin of *Peromyscus* mice expresses three different *Agouti* isoforms simultaneously, differing only in their first non-coding exon (1C, 1D, and 1E), the marked differences in overall *Agouti* expression seen between *P. maniculatus* strains (wideband vs. wild-type) and between *P. polionotus* subspecies (P. p. leucocephalus vs. *P*.*p. subgriseus*) are exclusively driven by one of the isoforms (1C). Thus, populations of *P. maniculatus* and *P. polionotus* experiencing selection pressures for light dorsal pigmentation have independently converged not only on the same gene, but also on the specific upregulation of the same isoform. One explanation for this result is that exon 1C has inherent sequence properties that result in large amount of protein (relative to other *Agouti* isoforms), indicating that such convergence in isoform upregulation may be driven by selection for the molecular mechanism promoting the highest amount of *Agouti* protein production. In support of this, we find that in a admixed population in the Sand Hills, dorsal color and *Agouti* gene expression is significantly associated with genetic variation around exon 1C and that this region shows a pattern of strong selection.

Given the patterns of phenotypic and gene expression association as well as the signatures of selection we have reported here and elsewhere (Linnen *et al*. 2013), it is likely that at least one causal polymorphism is located somewhere in the close vicinity of exon 1C (Fig. 5), as this population has low levels of linkage disequilibrium at the *Agouti* locus (Linnen *et al*. 2013). An important goal of future work is to characterize and functionally test the variants in this region, including indels and low-coverage SNPs that may have been absent in the capture-based genotype data. In the case of *P. polionotus* beach mice, despite the fact that selection on pigmentation is strong (*s* = 0.5; Vignieri *et al*. 2010) and *Agouti* is known to be a major contributor to pigment differences (Steiner *et al*. 2007; Mullen & Hoekstra 2008), any inferences of positive selection for regions in or around exon 1C are confounded by the unique demographic history of this species, which has experienced severe population bottlenecks associated with colonization events from the mainland to novel habitats in the Gulf Coast (Thornton *et al*. 2007; Domingues *et al*. 2012; Poh *et al*. 2014). Thus, it is not possible to evaluate with certainty whether specific regions in the *Agouti* locus show signatures of similar selective pressures between *P. maniculatus* and *P. polionotus*.

We find that the dorsal skin of *Peromyscus* expresses two isoforms that are not found in Mus or in other species (1D and 1E). However, it is unlikely that they have played a major role in the evolution of light coloration in *Peromyscus* because their expression patterns do not differ between the light and dark-colored strains examined here and do not follow the pulse of *Agouti* expression that occurs during hair growth (Vrieling *et al*. 1994). From a functional perspective, isoform 1D contains sequences that cause a marked repression of protein translation, so it is not surprising that this particular exon does not constitute a target for selection, and does not show differences in expression between *P. maniculatus* strains or *P. polionotus* subspecies. In the case of isoform 1E, our functional experiments suggest that the sequence of exon 1E found in wildtype *P. maniculatus* increases protein translation, whereas the exon 1E sequence found in wideband *P. maniculatus* does not impact translation. The two strains’ sequences differ only at the end of the exon, where wild-type *P. maniculatus* has a six base pair deletion (Fig S2, supporting information). Importantly, however, the exon 1E sequence found in an outgroup to the two strains, *P. maniculatus* rufinus, is identical to that found in wideband *P. maniculatus*, indicating that the deletion found in the wild-type 1E sequence is likely derived and arose after the split between wild-type and wideband populations. Thus, in the common ancestor of wildtype and wideband mice, exon 1C—and not exon 1E—would have been the only non-coding exon that promoted increased protein production. The driving forces underlying the evolution and maintenance of isoforms 1D and 1E in *Peromyscus* populations are yet unknown.

Since *Agouti*’s function is primarily linked to regulating pigment-type switching in melanocytes, changes affecting this gene are less likely to have negative pleiotropic consequences and thus, selection pressures for eliminating particular transcripts from populations may be relaxed. Alternatively, isoforms 1D and 1E could be playing a role in other aspects of *Agouti’s* function independent of its interaction with melanocytes that was not uncovered by our experiments, such as secretion and/or transport from the dermal papillae. Examining presence/absence and expression patterns of these isoforms in additional *Peromyscus* species and populations may shed light on some of these possibilities.

The findings presented here also bear on the molecular basis of convergent evolution. It has long been a topic of contention in evolutionary biology whether similar phenotypes that evolve independently tend to be generated by similar or different molecular changes (e.g., Stern 2013; Manceau *et al*. 2010; Rosenblum *et al*. 2010). Some studies of convergent phenotypes have found that they occur through independent mutations at different loci (e.g., Steiner *et al*. 2009; Weng *et al*. 2010; Kowalko *et al*. 2013), while others have found that similar evolutionary pressures on two populations can result in changes at the same gene (e.g., Woods *et al*. 2006), and in some cases, even the same amino acid substitutions (Zhen *et al*. 2012; van Ditmarsch *et al*. 2013). In the case of convergent pigmentation phenotypes in *Peromyscus*, not only is the same locus targeted, but the same pattern of isoform regulatory change has occurred—highlighting the various ways in which convergent evolution can occur at the molecular level.

Our results add an additional layer to the known mechanisms by which *Agouti* can play a role in the evolution of pigmentation phenotypes in *Peromyscus*. Cis-regulatory changes in *Agouti* are known to contribute to both the wideband phenotype in *P. maniculatus* (Linnen *et al*. 2009; 2013) and the beach mouse phenotype in *P. polionotus* (Steiner *et al*. 2007; Manceau *et al*. 2011). In addition, an amino acid change in the *Agouti* coding sequence of *P. maniculatus* wideband mice is strongly associated with light phenotypes and shows strong signatures of selection (Linnen *et al*. 2009; 2013). Here, we find that in addition to these changes, differences in *Agouti* isoform regulation have also been involved in the evolution of adaptive pigmentation variation in this genus. It is clear that alternative transcript processing provides a virtually limitless substrate for the generation of functional and structural transcriptomic and proteomic diversity, and genomic studies analyzing rates of alternative splicing and alternative promoter usage have revealed the importance of this mechanism in originating diversity at large taxonomic scales (Barbosa-Morais *et al*. 2012; Merkin *et al*. 2012). Our study, by providing an example that links alternative mRNA processing with adaptation to a known selective pressure in different subspecies between sister species, highlights the importance of this mechanism as a driver of diversification and adaptation at smaller taxonomic scales as well.

## Data accessibility

An Excel file containing the measurements for coat color, soil reflectance, and all gene expression values will be provided prior to publication.

## Acknowledgements

We thank Judy Chupasko and Mark Omura for help with specimen preparation, Catalina Perdomo for technical assistance with luciferase assays, and Nikki Hughes for providing logistical support. RM and the molecular work was supported by a Swiss National Science Foundation grant to HEH. TAL was supported by a Herchel Smith-Harvard Undergraduate Research Fellowship; CRL was supported by a Ruth Kirschstein National Research Service Award from NIH; HEH is an Investigator of the Howard Hughes Medical Institute. This work was supported by a grant from the Swiss National Science Foundation.

## Author contributions

RM, CRL, and HEH conceived the project and designed the experiments. RM, TAL, and CRL performed experiments and analyzed data. All authors wrote the paper.

